# Game theoretic methods in population dynamics

**DOI:** 10.1101/056341

**Authors:** David Liao

## Abstract

Game theoretic methods are used to model the dynamics of cellular populations (tissues) in which it is necessary to account for changes in cellular phenotype (usually proliferation) resulting from cell-cell interaction. Results include prediction of long-term steady-state “equilibria” and transient dynamics. These results can be useful for predicting relapse after cytoreduction, assessing the efficacy of alternating combination therapy, and interpreting biopsy specimens obtained from spatially heterogeneous tissues. Mathematical tools range from simple systems of differential equations to computational techniques (individual-based models).

## I. PROBLEMS IN CANCER

One of the goals of research into basic cancer biology and cancer therapy is to understand how the current condition of a population of cells can be used to predict future dynamics of the tissue, and ways to modify outcomes through therapeutic intervention. Addressing this goal promises to aid the search for prognostic markers to stratify risks for future occurrence and recurrence, the modeling of risks of developing drug resistance, the understanding of delayed recurrence after dormancy, the prediction and control of metastasis, and the development of therapeutics and dosing schedules. It is increasingly recognized that analyses of intracellular genetic regulatory networks will not alone be sufficient to provide theoretical insights for fully addressing these goals. Given the sophisticated interactions between cell types in both normal and malignant tissue microenvironments [1,2], mathematical modeling of the influences that subpopulations of cells (and non-cellular materials) exert on cellular phenotypes will be required. “Game theoretic” methods refer to a broad range of modeling techniques that incorporate these influences, often by describing how the fractional proliferation rates of cellular subpopulations vary with the fractional representation of these subpopulations in the population overall (Fig. 1).

**Figure 1.**
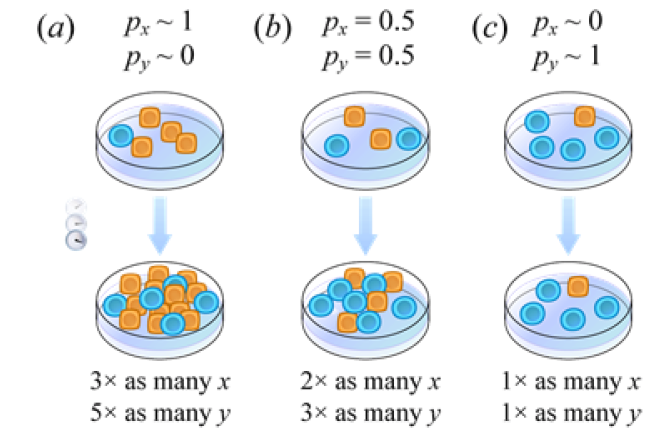
Expansion factor of a subpopulation depends on population composition. Reprinted from [5] Fig. 2.

## II. ILLUSTRATIVE RESULTS OF APPLICATION OF METHODS

Game theory has been used to understand dynamic changes in the proportions of cellular subpopulations during disease progression, treatment, and relapse. According to a mathematical model of interactions between multiple myeloma, osteoblast, and osteoclast cells developed by Dingli *et al.* [3], partial cytoreduction of the multiple myeloma subpopulation fails to eliminate the stable equilibrium point associated with eventually re-established dominance of the multiple myeloma subpopulation (Fig. 2). In addition to predicting equilibria, game theory can be used to predict transient dynamics preceding long-term steady states. For example, Basanta *et al.* [4] developed a model of glioblastoma multiforme that predicted that eventual dominance by the invasive phenotype might be preceded by oscillations in population composition.

**Figure 2.**
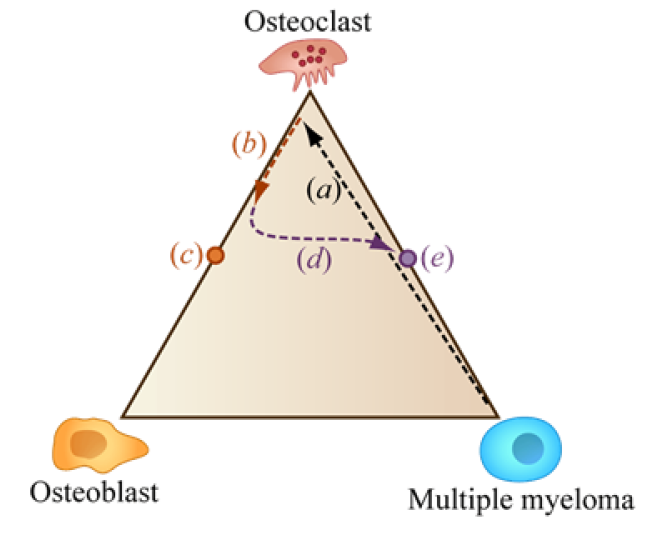
Using a simplex to illustrate eventual regrowth of multiple myeloma following therapeutic cytoreduction. Adapted by permission from Macmillan Publishers Ltd on behalf of Cancer Research UK: *<u>Br. J. Cancer.</u>* [3], Fig. 4(e), copyright 2009.

Velocity fields provide a graphical representation of replicator dynamics equations. In two-dimensions, such an illustration on a coordinate grid is referred to as a phase portrait (“simplex” in three dimensions). Wu *et al.* used phase portrait analysis to illustrate how multiple myeloma and stromal cells treated with doxorubicin in microfabricated habitat structures can initially exhibit an increase in the multiple myeloma subpopulation before eventually becoming dominated by stroma [6]. For such a trajectory, apparent failure can be merely temporary. Liao *et al.* [5] proposed using velocity fields to evaluate combination therapies. The efficacy of alternating between two therapies might be predicted by measuring the angles formed by quivers on a phase portrait (Fig. 3).

Game theory has also been applied to model dynamics according to which a treated tumor and oncologist respond to each other [7].

**Figure 3.**
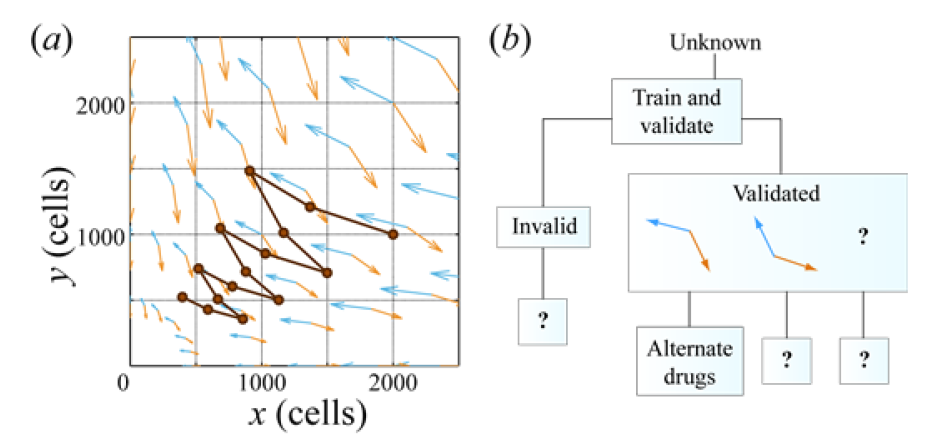
Game theory can be used to evaluate candidate schedules for combination therapy. Adapted from [5] Fig. 4.

Kaznatcheev *et al.* [8] incorporated spatial effects into an analysis of the “go vs. grow” game to illustrate why core biopsies need to be interpreted with caution. In Fig. 4, cells in the interior bulk of a tissue interact with more neighbors than cells near a fixed edge (e.g. tissue interface). The number of neighbors in a cellular network strongly influences which phenotype will dominate (see Ohtsuki-Nowak transform below). Histology supports concern that the invasive phenotype might dominate only a thin tissue layer near the interface. Thus, dominance of the invasive phenotype near the tissue interface might not be apparent from a core biopsy that mainly samples the bulk, diluting the contribution of cells near the edge.

**Figure 4.**
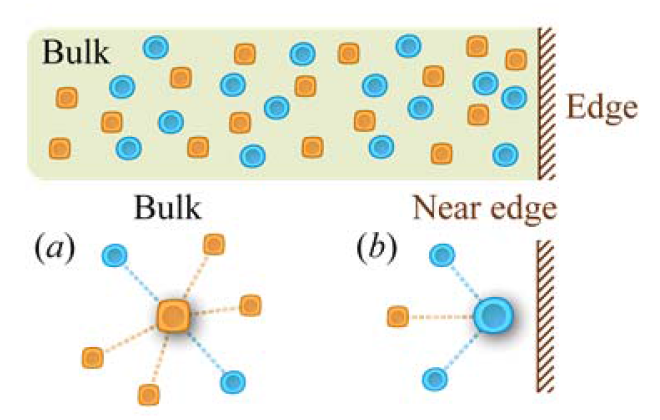
Biopsy with poor spatial resolution can provide misleading population composition information.

Based on recent studies in microbial populations, we anticipate future applications of evolutionary game theory and related ecological approaches to cancer research. For example, the phenomenon of “critical slowing down” that can predict imminent population collapse [9,10,11] might be applied to detect when normal or malignant cell populations are near a critical transition.

## III. QUICK GUIDE TO METHODS

We highlight two types of game theoretic models. For additional reading, tutorials [5] and [12] are suggested for mathematical novices. For a brief overview of methods see [13], and for more mathematical reviews, see [14] and [15] and additional reviews listed in [8].

### A. Well-mixed, continuous-time replicator dynamics

One of the simplest forms of evolutionary game theory commonly applied can be derived from assumptions that cells are (1) vigorously mixed so that each cell collides with cells of various subtypes at frequencies proportional to the global population fractions of those cellular subtypes and that (2) cell-cell contact events temporarily modify rates of proliferation (Fig. 5) in a linear way.

**Figure 5.**
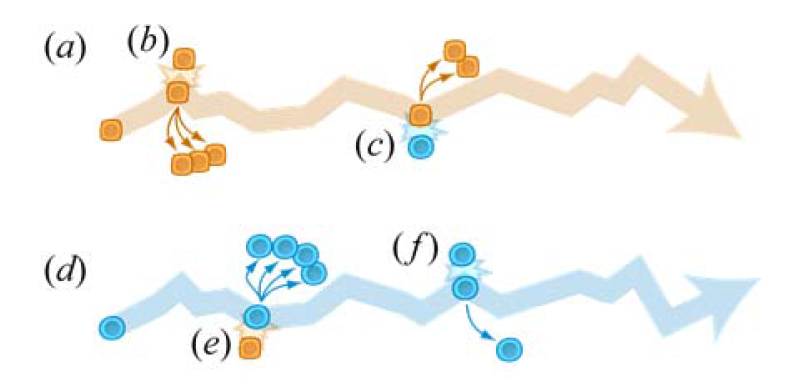
Assumption that pairwise collisions between cells induce proliferation events. Adapted from [12] Fig. 2.

Equations 1 below follow from these assumptions.

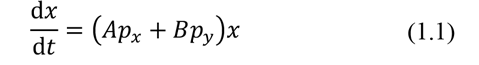

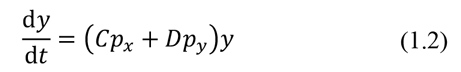

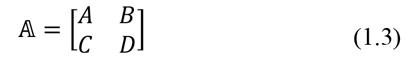

Here, time rates of change (d/d*t*) of subpopulations (e.g. numbers of cells of type *x* and *y*) are proportional both to the cell numbers *x* and *y*, and to fitnesses that are linear functions of the ***global*** population fractions, *p_x_* and *p_y_*, of the cell types *x* and *y*, respectively. For example, in Eqn. 1.1, coefficient *A* is the replicative fitness of cells of type *x* in a population consisting only of cells of type *x*, coefficient *B* is the replicative fitness of rare cells of type *x* in a population consisting almost completely of cells of type *y*, and the fitness function *Ap_x_* + *Bp_y_* linearly interpolates between these extremes to provide fitness values for intermediate population compositions. Taken together, as in Eqn. 1.3, the coefficients are often referred to as a payoff matrix.

### B. Ohtsuki-Nowak transform

The preceding model can be made more realistic by assuming that cells interact with local neighbors and proliferate on a spatially-resolved random network. Fig. 6 illustrates examples of compositions of local neighborhoods that can be realized by a spatially-resolved model, but not by a well-mixed model. In a well-mixed population, cells at different locations, e.g. (*a*), (*b*), and (*c*) confront local neighborhoods with the same demographic compositions. In a less thoroughly mixed population, a cell might find itself in the midst of (*d*) a neighborhood similar to those expected for a well-mixed population, (*e*) a neighborhood with greater homogeneity than expected for a well-mixed population, e.g. the cell is surrounded by more cells of its own type than expected on average, or (*f*) a neighborhood with greater heterogeneity than expected for a well-mixed population, i.e. a neighborhood in which heterotypic interactions are more common than expected on average. Panels (*e*) and (*f*) illustrate two scenarios that can never happen in a well-mixed population and that require that the original payoff matrix, 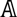, in Eqn. 1.3 be replaced by a transformed matrix in order for Eqns. 1.1 and 1.2 to correctly describe the global dynamics of the spatially-resolved network.

**Figure 6.**
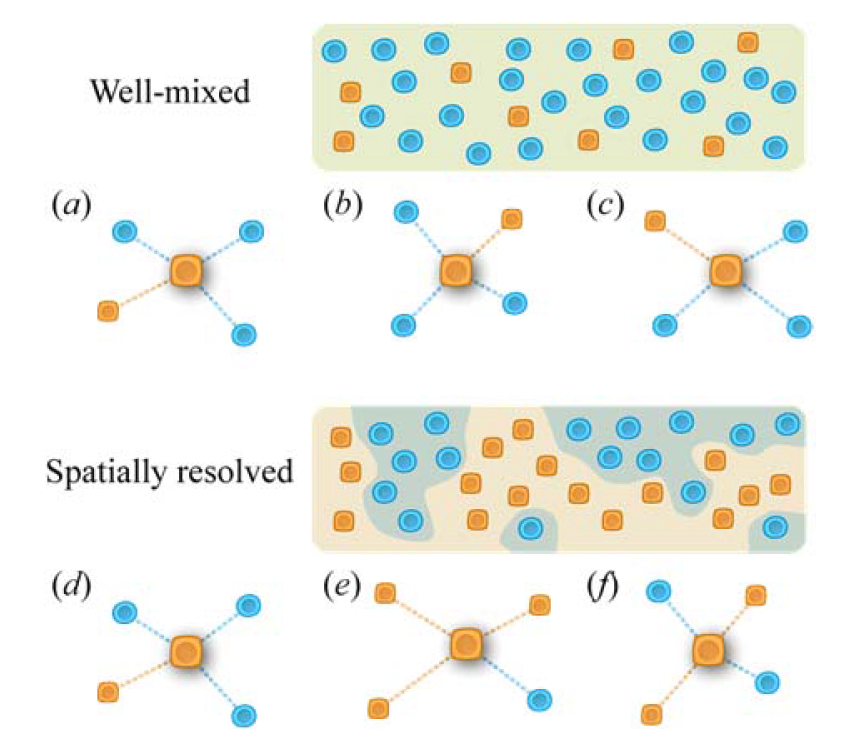
Heuristic for explaining terms in Ohtsuki-Nowak transform.

The “transformed” matrix, *ON_k_*(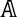), in Eqn. 2, is obtained using the Ohtsuki-Nowak transformation [8,16,17]. This transformation relies on assuming that statistical correlations between cells can be neglected for non-nearest neighbors.

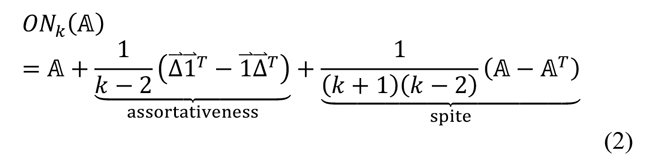

The exact form of Eqn. 2 varies with details of the particular birth-death update process under investigation, but the general form has three terms (original matrix accompanied by two correction terms, as heuristically anticipated from the scenarios in panels (e) and (f) in Fig. 6). The number of neighbors with which an individual immediately interacts is *k*. One way to remember that the dynamics (including steady state population compositions) depends on the neighborhood size is to remember that a cell type that is easily outcompeted in a well-mixed population might survive in a less thoroughly mixed population when protected by being locally surrounded by other cells that do not outcompete it.

The Ohtsuki-Nowak transform was the technique that enabled the analysis of edge effects and their consequences for biopsy analyses, as described in [8].

### C. Individual-based models

While the Ohtsuki-Nowak transform analytically describes global population dynamics for large random networks with local cell interactions, computational techniques are typically used to model detailed histories of individual cells. Example capabilities of individual-based models (IBMs, agent-based models, ABMs) are described elsewhere in this guide [18]. In IBMs, individual cells carry out scripts that prescribe phenotypes to be expressed (e.g. proliferation, movement, changes in cell type) according to the local environments in which individual cells are found.

### D. Scales at which these methods are useful

While examples in this summary relate to interactions between cells, the same mathematical techniques can be applied at larger scales to describe interactions between tissues and at smaller scales to describe interactions between pools of molecules in intracellular regulatory networks. Insights into network topology at subcellular scales might provide a head start for understanding topologies of cell-cell interactions that have been evolutionarily shaped.

## ONLINE RESOURCES

a. Kaznatcheev A. *Theory, Evolution, and Games Group*. Blog: <http://egtheory.wordpress.com/>
b. Kaznatcheev A, Scott J, Basanta D. *Evolutionary Game Theory.* An excellent Google community for questions and discussions. <http://plus.google.com/u/0/communities/114797456234998377606>
c. Liao D. Tour: Game theory. Educational materials for tutorials in *Interface Focus.* <http://quant.bio/tour.egt.php>

## Notes

Research was supported by award U54CA143803 from the US National Cancer Institute. The content is solely the responsibility of the authors and does not necessarily represent the official views of the US National Cancer Institute or the US National Institutes of Health.

